# Brain-to-brain synchrony between students and teachers predicts learning outcomes

**DOI:** 10.1101/644047

**Authors:** Ido Davidesco, Emma Laurent, Henry Valk, Tessa West, Suzanne Dikker, Catherine Milne, David Poeppel

## Abstract

Little is known about the brain mechanisms that underpin how humans learn while interacting with one another in ecologically-valid environments (1-3). This is because cognitive neuroscientists typically measure one participant at a time in a highly constrained environment (e.g., inside a brain scanner). In the past few years, researchers have begun comparing brain responses across individuals (4-6) demonstrating that brain-to-brain synchrony can predict subsequent memory retention (7-9). Yet previous research has been constrained to non-interacting individuals. Surprisingly, the one study that was conducted in a group setting found that brain synchrony between students in a classroom predicted how engaged the students were, but not how much information they retained (10). This is unexpected because brain-to-brain synchrony is hypothesized to be driven, at least partially, by shared attention (11, 12), and shared attention has been shown to affect subsequent memory (13). Here we used EEG to simultaneously record brain activity from groups of four students and a teacher in a simulated classroom to investigate whether brain-to-brain synchrony, both between students and between the students and the teacher, can predict learning outcomes (Fig. 1A). We found that brain-to-brain synchrony in the Alpha band (8-12Hz) predicted students’ delayed memory retention. Further, moment-to-moment variation in alpha-band brain-to-brain synchrony discriminated between content that was retained or forgotten. Whereas student-to-student brain synchrony best predicted delayed memory retention at a zero time lag, student-to-teacher brain synchrony best predicted memory retention when adjusting for a ∼200 millisecond lag in the students’ brain activity relative to the teacher’s brain activity. These findings provide key new evidence for the importance of brain data collected simultaneously from groups of individuals in ecologically-valid settings.

**Highlights:**

- Electroencephalogram (EEG) was concurrently recorded in a simulated classroom from groups of four students and a teacher.
- Alpha-band (8-12Hz) brain-to-brain synchrony predicted students’ performance in a delayed post-test.
- Moment-to-moment variation in alpha-band brain-to-brain synchrony indicated what specific information was retained by students.
- Whereas student-to-student brain synchrony best predicted learning at a zero time lag, student-to-teacher brain synchrony best predicted learning when adjusting for a ∼200 millisecond lag in the students’ brain activity relative to the teacher’s brain activity.

## Results and Discussion

### Behavioral results

Students’ content knowledge was assessed a week before the EEG session, immediately following each one of four mini-lectures, and one week later (Fig. 1B; See Methods). As expected, students’ content knowledge significantly increased from the pre-test (0.43±0.02; mean ± standard deviation of the mean) to the immediate post-test (0.73±0.02; F(1,30)=210.76; p<10^−13^), and from the pre-test to the delayed post-test (0.64±0.02; F(1,30)=93.48; p<10^−10^; Fig. 2A). The retention of content knowledge significantly declined over the course of the week between the immediate and delayed post-tests (F(1,30)=46.00; p<10^−6^). The difference between the pre- and delayed post-test scores was considered as the outcome variable in all subsequent analyses.

**Figure 1.**
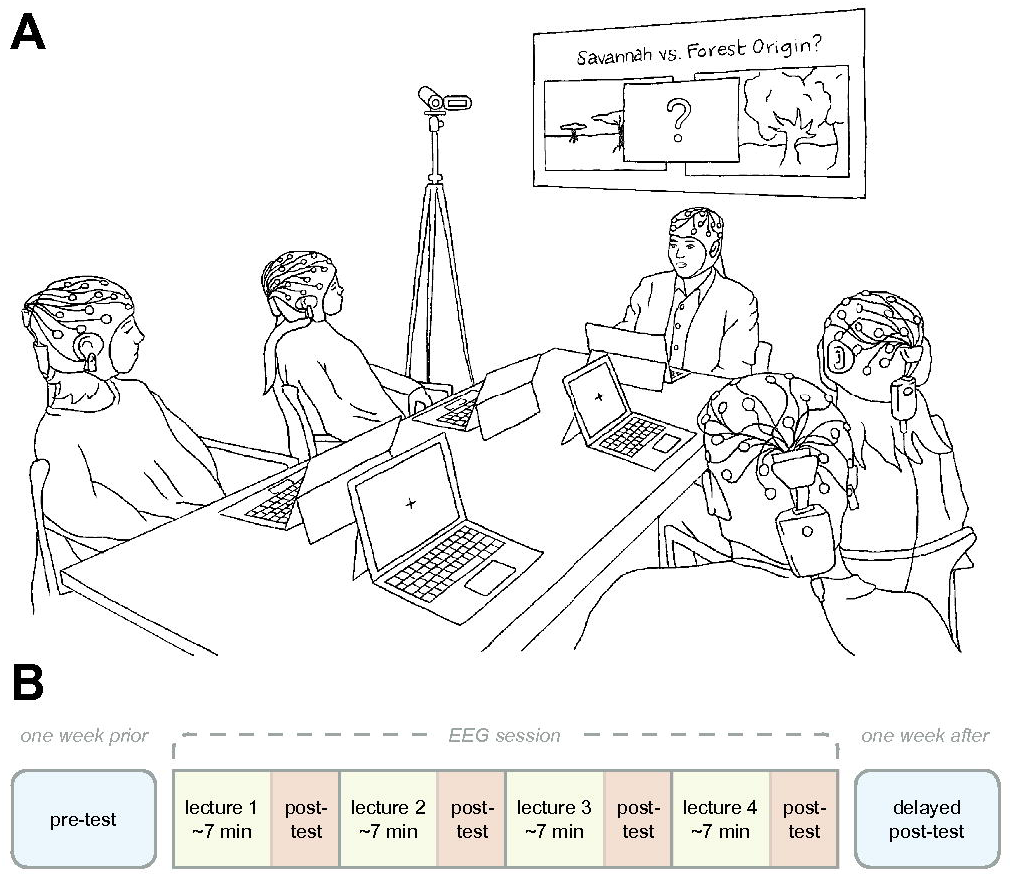
Experimental setup and timeline. (A) Four students and a teacher were concurrently measured with EEG during a science lesson; (B) The lesson comprised four mini-lectures, each followed by a post-test. Pre-test and delayed post-tests were administered one week prior to and one week following the EEG recording session.

**Figure 2.**
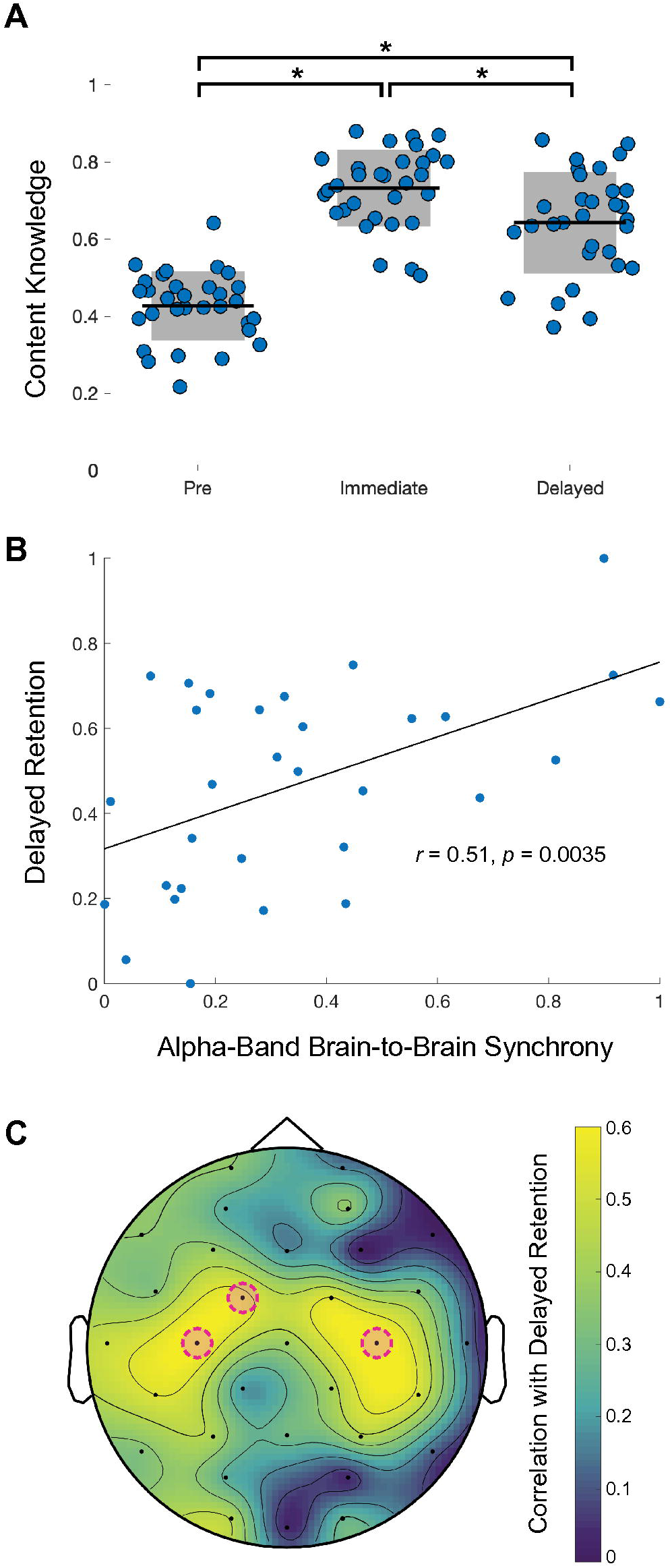
Brain-to-brain synchrony predicts delayed memory retention. (A) Proportion of correct answers (content knowledge) for the pre-test, immediate post-, and delayed post-tests. Each dot corresponds to one participant, horizontal black lines depict the mean for all students; grey regions represent one standard deviation; (B) Alpha-band student-to-student brain synchrony significantly predicted delayed retention; All values were normalized to a 0–1 scale (max-min) for presentation purposes, and each dot reflects the brain synchrony (quantified by CCorr) between one student and all the other students in the group, averaged across the four lectures; (C) Spatial distribution of the relationship between delayed retention and alpha-band brain-to-brain synchrony. Electrodes circled in pink were found to significantly predict delayed retention (p<0.05; FDR corrected).

### Student-to-student brain synchrony and memory retention

For each student dyad in a given group, Circular Correlation values (CCorr; (14)) were computed for all one-to-one paired combinations of electrodes (e.g. O1-O1, P3-P3 etc.; (10, 11)) (Fig. S1 and Methods). We first assessed whether student-to-student brain synchrony can predict memory retention. Since students were nested within groups, we constructed a multilevel model wherein brain-to-brain synchrony across three frequency bands (Theta: 3-7Hz; Alpha: 8-12Hz; Beta: 13-20Hz) were considered as level 1 predictors (see Methods). This analysis revealed that delayed memory retention was significantly predicted only by alpha-band brain-to-brain synchrony (Alpha: F(1,8.35)=8.76; p=0.017; Theta: F(1,17.29)=0.01; p=0.92; Beta: F(1,26.60)=1.34; p=0.26; Fig. 2B and S2). This finding is consistent with prior research, which links alpha oscillations with attention (15-19). The alpha-band has also been shown to be the most robust frequency range for brain-to-brain synchrony (20). Therefore, all subsequent analyses were focused on the Alpha band.

Next, rather than computing brain-to-brain synchrony across all 32 channel pairs, we compared three regions of interest (ROIs): posterior, central, and frontal (see Methods). Students’ delayed memory retention was significantly predicted only by the central ROI (F(1,25.35)=9.45; p=0.005), not by the posterior (F(1,26.96)=2.13, p=0.16) or frontal ROIs (F(1,25.91)=0.23, p=0.64) (Fig. S3). Furthermore, alpha-band brain synchrony significantly predicted delayed retention at the individual electrode pair level. Three of the 32 electrode pairs (C3-C3, C4-C4 and FC1-FC1) significantly predicted delayed retention (p<0.05; false-discovery-rate (FDR) corrected; Fig. 2C).

### Moment-to-moment variations in brain synchrony predict delayed retention

Typically, brain-to-brain synchrony is computed over an extended period of time (e.g. the entire duration of a lecture or a video (7, 10, 11)). In order to examine whether moment-to-moment variations in brain synchrony could indicate what specific information students retained a week later, we transcribed all the lectures and identified *when* the teacher provided information to answer each one of the test questions (Fig. 3A). For each student, a question was either classified as (1) “learned” - if the student answered it correctly in the delayed post-test, but not in the pre-test - or (2) “not learned” - if a student’s answer had not changed between the pre- and the post-tests (see Methods). Brain synchrony was then averaged separately for lecture epochs associated with learned and not learned questions. Across the three ROIs that were examined, only brain synchrony within central electrodes was significantly higher for learned compared to non-learned information (learned: 1.09±0.04; not learned: 0.97±0.03; F(1,38.69)=8.85; p=0.005; Fig. 3B).

**Figure 3.**
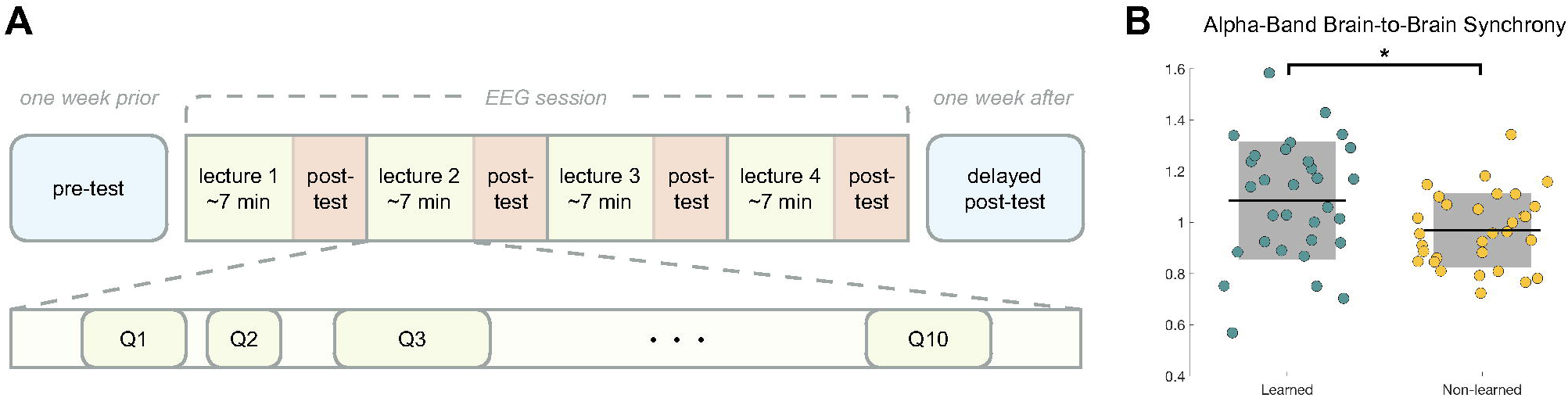
Moment-to-moment variations in alpha-band brain-to-brain synchrony and learning. (A) Question-specific time intervals where relevant content was delivered by the teacher were identified based on the lecture transcript; (B) Moment-to-moment variations in alpha-band brain-to-brain synchrony across central electrodes significantly discriminated between information that was learned and not learned at the individual question level. For each student, brain synchrony was normalized by dividing the epoch-by-epoch CCorr values by the averaged CCorr value across the entire duration of the lecture.

### Student-to-teacher brain synchrony

So far, we have only considered brain-to-brain synchrony between students rather than between the students and the teacher. As students only listened to the lectures, we hypothesized that student-to-student brain synchrony would best predict delayed retention at a zero lag (i.e. when students’ brain activity is concomitantly aligned). In contrast, because the teacher served as the speaker and the students as listeners, we expected student-to-teacher brain synchrony would best predict delayed retention at a non-zero lag (i.e. when the teacher’s brain activity is shifted backwards relative to the student’s brain activity) (21). Similar to (22), we computed time-lagged student-to-student and student-to-teacher brain synchrony and then correlated brain synchrony with delayed retention for each time lag (see Methods). On average, the correlation between student-to-student synchrony and delayed retention indeed peaked for zero-lagged synchrony (Fig. 4A). The correlation between student-to-teacher synchrony and delayed retention, on the other hand, showed two peaks: a peak at around -200 msec lag between the student and the teacher (i.e. teacher’s brain activity preceding students’ by about 200 msec) and a peak at around +250 msec lag (i.e. students’ brain activity preceding the teacher’s by about 250 msec) (Fig. 4B). To better understand this result, we plotted the lag at which the correlation between student-teacher brain synchrony and memory retention was maximal for each electrode pair (Fig. 4C). Intriguingly, while most electrode pair correlations peaked when the teacher’s brain activity preceded that of the students, central and frontal electrodes showed the reverse pattern, where the correlation between student-to-teacher synchrony and delayed retention peaked when the student’s brain activity preceded the teacher (Fig. 4C). This finding is consistent with previous research on speaker-listener brain synchrony, which has demonstrated that listeners’ brain activity is coupled with speakers’ at a delay (21, 23). However, these previous studies used methods with low temporal resolution (functional Magnetic Resonance Imaging (fMRI) and functional Near-Infrared Spectroscopy (fNIRS)), and thus could not accurately estimate the speaker-listener delay. A delay of roughly 200 msec is consistent with the time scale of speech processing (24). Our finding that in central and frontal electrode pairs, the correlation between student-to-teacher synchrony and delayed retention peaked when the student’s brain activity preceded the teacher is consistent with previous fMRI research (21, 25), and might reflect predictive anticipation of upcoming input by students (26).

**Figure 4.**
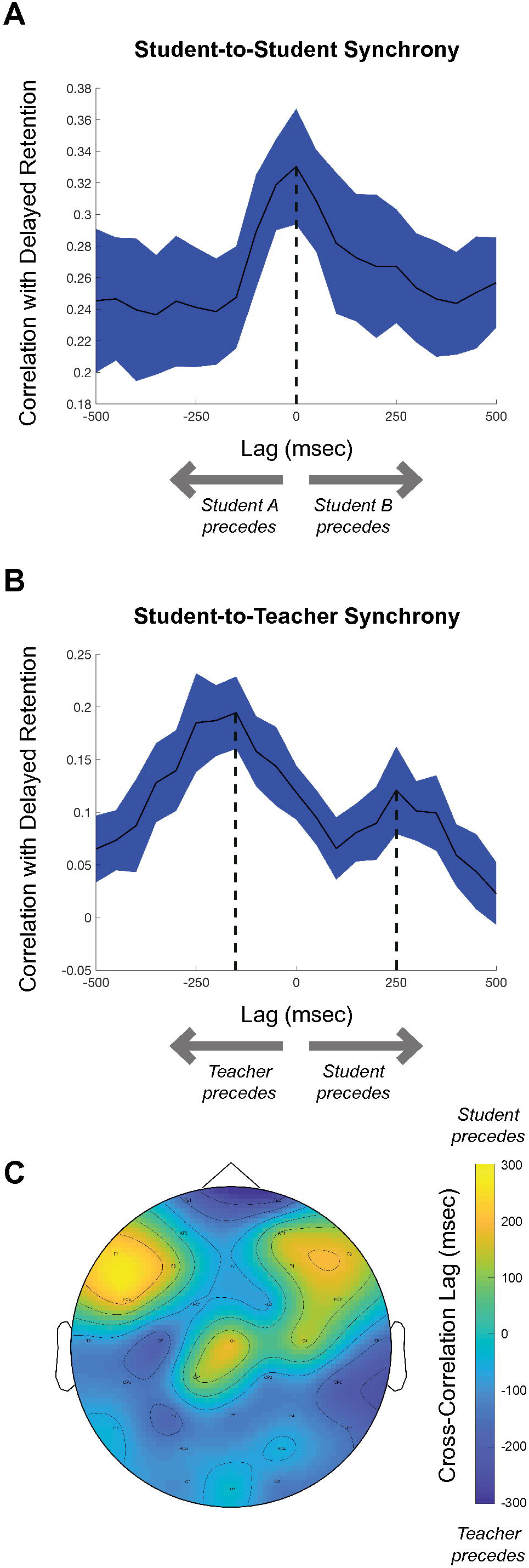
Time-lagged cross-correlation between brain-to-brain synchrony and delayed retention. (A) Cross-correlation between student-to-student brain synchrony and delayed retention as a function of temporal lag between students’ brain activity; (B) Cross-correlation between student-to-teacher brain synchrony and delayed retention as a function of temporal lag between the student’s and teacher’s brain activity. For both (A) and (B), cross-correlation was computed for each electrode pair and then averaged across pairs (total of 32 pairs). (C) Spatial distribution of the temporal lag that produced the highest correlation between student-to-teacher brain synchrony and delayed retention. Electrodes are color coded by temporal lag: student precedes (yellow) to teacher precedes (blue).

The aim of the current study was to explore whether brain synchrony across students and teachers can predict long-term memory retention. While brain-to-brain synchrony has been associated with several classroom-related variables (for example, students’ engagement and social closeness; (10, 11, 27-29)), there are conflicting results about its relationship with learning outcomes. Bevilacqua et al. (10) collected EEG data from a group of 12 high school students and their teacher during regular biology lessons. Student-to-teacher brain synchrony predicted how engaged students were and how close they felt toward the teacher, but it was not significantly associated with how well students retained class content. Cohen et al. (7) measured brain-to-brain synchrony between students who watched instructional videos individually. Even though the students were not measured concurrently, brain-to-brain synchrony was found to predict their immediate memory retention (7). These contradicting findings could be explained by methodological differences. While Bevilacqua et al. (10) used commercial-grade EEG devices in a classroom environment, Cohen et al. (7) used research-grade EEG devices in a lab setting. Our findings are consistent with several other previous studies that were conducted outside of the education context. Hasson et al. (9), using fMRI, demonstrated that inter-subject correlations during movie viewing correlate with delayed memory retention. Further, Cohen & Parra (8) reported that inter-subject correlations of broadband EEG signals during the presentation of auditory narratives predicted subsequent memory of the narratives. Critically, however, in all these previous studies (7-9), participants were measured individually, and thus there was no social interaction between participants during stimulus encoding. In the absence of such interaction, previous studies were not able to assess whether synchrony between students and teachers is related to learning outcomes. The current study substantially extends previous research by demonstrating that both student-to-student and student-to-teacher alpha-band brain synchrony are associated with long-term memory retention. This effect was observed both at the whole brain level (Fig. 2B) and for individual electrode pairs (Fig. 2C), but was constrained to the alpha-band (Fig. S2). Furthermore, moment-to-moment variations in brain synchrony during lectures was shown to significantly discriminate between retained and non-retained information (Fig. 3).

While the phenomenon of brain-to-brain synchrony is not yet fully understood, Dikker et al. (11) proposed that shared attention plays a crucial role. At the most basic level, brain-to-brain synchrony is driven by stimulus entrainment: as all students are presented with the same input (e.g. teacher’s voice), their brain activity becomes entrained to that stimuli. Critically, since stimulus entrainment is modulated by attention (30, 31), brain-to-brain synchrony increases when students are engaged in a task and decreases when students disengage (7, 10, 11). The hypothesis that brain-to-brain synchrony is partially driven by shared attention is consistent with the current study: when students pay attention to information provided by a teacher, their brain synchrony with the teacher and other students increases, as does their tendency to retain information. The current study focused on brain-to-brain synchrony in the alpha band (8-12 Hz) since there is extensive research linking the alpha rhythm to attention. While traditionally associated with cortical idling (32), it is currently thought that the alpha rhythm is involved in actively suppressing task-irrelevant processing (15-19). Further, there is substantial evidence that the phase and amplitude of alpha-band oscillations prior to and during stimulus presentation influences subsequent stimulus processing (33, 34).

Further research, possibly in more controlled experiments, is needed to understand the neural dynamics that give rise to brain-to-brain synchrony. Future research might examine not only what conditions enhance brain synchrony, but also under what circumstances brain synchrony is diminished, and what the behavioral consequences of decreased neural synchrony are. It should go without saying that the methods we have to study the human brain do not permit more neurobiologically granular, mechanistic characterization. That being said, the measures that we have used here yield unanticipated new insights into how learning in a group context is reflected in the brain dynamics of teachers and learners.

## Acknowledgments

We thank U. Hasson for comments on an earlier version of this manuscript, and O. Dagan, E. Theisen, D. Bevilacqua, G. Ali, and S. Azeka for their assistance in data collection and data preprocessing. This study has been supported by NSF grant #1661016.

## Author Contributions

Conceptualization, I.D., H.V., S.D., C.M., and D.P.; Investigation, I.D., E.L., and H.V.; Writing, I.D., H.V., S.D., T.W., and D.P.; Supervision, D.P.

## Methods

### Participants

42 participants (28 females) were recruited and measured in groups of three or four students. All participants satisfied the following criteria: (i) native English speaker; (ii) right hand dominant; (iii) between the ages of 18 and 30 (mean age: 20.6; s.d.: 3.0 years); (iv) non-science major, if applicable; (v) no known history of neurological abnormalities. All participants completed high school, with the majority (76.2%) being current college undergraduates. All participants provided written informed consent, and the experimental protocol was approved by the Institutional Review Board of New York University. In two of the 11 groups, due to technical issues, only two participants had usable EEG data; as student-to-student brain synchrony could not be determined for all dyads, all participants within these two sessions were excluded from analysis (N=7). Four additional subjects were omitted from analysis: two due to poor quality EEG data and the other two since they scored higher in the pre-than the post-test for the majority of lectures (i.e. they did not demonstrate any learning gains). Thus, the final sample consisted of 31 participants (21 female).

The two teachers (1 female) were professional high school science teachers. The female teacher led four sessions, and the male teacher led five of the nine lessons included in analysis. The teachers had no prior acquaintance with the students.

### Procedure

Students were seated evenly and randomly around a table, and the teacher was seated at the head of the table (Fig. 1). The experiment took place in a laboratory classroom equipped with a projector and three video cameras. Following EEG set up, baseline EEG recordings (eyes-open and eyes-closed) were taken to test data quality. The lesson comprised of four teacher-led lectures (6:43.26±0:45.93 minutes-long; mean±s.d.) on discrete topics in Biology and Chemistry: Bipedalism, Insulin, Habitats and Niches, and Lipids. Slides were projected onto a screen behind the teacher and controlled by the teacher via a tablet computer (see Fig. 1). In order to minimize speaking- and movement-related artifacts, students were instructed to sit still, minimize head motion, and refrain from asking questions during the lectures. Each lecture was preceded by either no activity or one of three brief pre-lecture activities, where students could interact more freely with one another and with the teacher: a discussion-based activity, a short quiz where students answered three topic-related questions and then observed the distribution of answers across their group, or a short video related to the lecture topic. Activity–lecture combinations and order were randomly pre-assigned and counterbalanced across groups. Each lecture was immediately followed by a brief topic-specific assessment to gauge lecture engagement and content knowledge (see below). Assessments were administered via a tablet computer that was placed next to each student. The lesson concluded with one final three-minute eyes-open baseline recording. The same four content knowledge assessments were given to participants individually both one week prior to (pre-test) and one week following (delayed post-test) his or her corresponding group session (Fig. 1B).

### EEG hardware and data collection

Participants’ EEG activity was recorded using a 32-channel Neuroelectrics Enobio 32 5G gel sensor system (sampling rate: 500Hz). A dual earclip electrode served as a common unipolar reference. Electrode placement followed the standard 10-10 EEG system. The Neuroelectrics Instrument Controller (NIC2) software application was used to record data and assess signal quality. Data was aligned between students and the teacher post-hoc at the millisecond level using wireless triggers that were sent every second by a tablet computer via Lab Streaming Layer (LSL; (35)).

### Quantifying memory retention

For each lecture, memory retention was measured using 10 multiple-choice questions and one short answer question (only the multiple-choice questions were used in the current analysis). Questions were developed by the two participating teachers and reviewed by an independent education specialist (see Table S1). In order to measure changes in content knowledge at the individual question level, the same content questions were used in the pre-test, immediate, and delayed post-test. Note that in order to minimize priming effects, the pre-test was administered a week before the EEG session (Fig. 1B). Students’ scores ranged between 0 to 100 (% of multiple choice questions that were answered correctly). The difference in student scores between the pre-test and delayed post-test were averaged across lectures and used as the main outcome variable throughout this study. Note that the participants were instructed not to discuss the material with each other or read about the topics covered in the lectures between study sessions.

All the lectures were audio recorded and synced to the EEG data via LSL. The lectures were transcribed and for each content question, time intervals in which information necessary to answer the question were identified. This enabled matching question-specific EEG data with students’ answers to these questions (Fig. 3).

### EEG Preprocessing

All preprocessing was carried out using Matlab R2018b in conjunction with EEGLAB 14.1.1b (36). Only data recorded during lecture presentation was included in the analysis. After band-pass filtering (0.5 to 35 Hz), noisy channels were identified and removed using a combination of automatic channel rejection (kurtosis, z-score=3) and inspection of channel power spectra. Continuous EEG data were then epoched into 1-second intervals and visually inspected for non-neural artifacts. Independent component analysis (ICA) was then conducted to identify and remove components that were associated with eye blinks and eye movements (37). Finally, abnormal residual epochs with signals outside of -100 to 100 µV range were automatically tagged and visually inspected. It should be noted that due to the nature of this experiment, teacher data were inherently noisier than those of students. As a result, a more stringent data removal approach was required to obtain high-quality teacher data (See Table S2).

### EEG analysis

The data were analyzed using custom-built Matlab code and the FieldTrip toolbox (38). Following preprocessing, EEG data were filtered between 8- and 12-Hz using Butterworth filters of order four, and Hilbert transform was used to compute the instantaneous phase. For each 1-second epoch and for each one-on-one paired combinations of electrodes (e.g. O1-O1, P3-P3 etc.) (10, 11), CCorr (14) was calculated by:

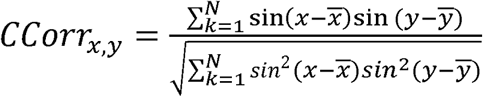

Where x corresponds to an EEG channel in participant #1 and y corresponds to the same EEG channel in participant #2. 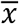 and 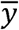 are the mean directions of the EEG channels. N is the number of samples in each epoch (500).

In the calculation of CCorr, only overlapping EEG channels and epochs were considered. In other words, if a specific channel or epoch was excluded in participant #1, they were also excluded in participant #2 (10, 11).

CCorr was chosen because it has been shown to be the least sensitive to spurious couplings of EEG hyperscanning data (39). CCorr values were calculated for each pair of students within a group and between each student and the teacher (see Fig. S1). Calculated CCorr values were normalized by Fisher’s Z transformation and averaged across epochs, lectures, and electrode pairs.

Moment-to-moment analysis (Fig. 3): In this analysis, rather than averaging CCorr values across the entire duration of each lecture, data were averaged across question-specific epochs identified based on the lecture transcript (Fig. 3A). Since information needed to answer a specific question could have been mentioned more than once in the lecture, all these instances were included in the analysis. However, to obtain stable brain synchrony estimates, only lecture-segments of 3 seconds or longer were included. A question was categorized as “learned” if a student answered it correctly in the delayed post-test, but not in the pre-test. A question was categorized as “not learned” if a student’s answer has not changed between the pre- and the delayed post-test (i.e. the student either already knew the answer to the question before the lecture, or answered it correctly before the lecture and incorrectly after the lecture). For each student, brain synchrony was normalized by dividing the epoch-by-epoch CCorr values by the averaged CCorr value across the entire duration of the lecture.

Time-lagged cross-correlation analysis (Fig. 4): Similar to (22), for each student-student dyad, the time course of one of the students was shifted either backward or forward in the range of -500 msec to +500 msec in steps of 50msec. Similarly, for each student-teacher dyad, the time course of the teacher was shifted between -500 msec to +500 msec in steps of 50 msec with respect to the time course of the student. Brain synchrony was computed for each temporal lag and then correlated with delayed retention. This was done for each electrode pair and then the Fisher’s Z transformed correlation values were averaged across electrode pairs (Fig. 4A-B).

### Statistical analysis

Since students were nested within groups, data were analyzed using multilevel modeling treating group as the unit of analysis to control for nonindependence in student responses. The MIXED procedure in SPSS was used. Alpha-band brain-to-brain synchrony was included as a level 1 predictor. Delayed retention was treated as the outcome variable.

