## Supporting Information for "Brain-to-brain synchrony between students and teachers predicts learning outcomes"

Davidesco et al.

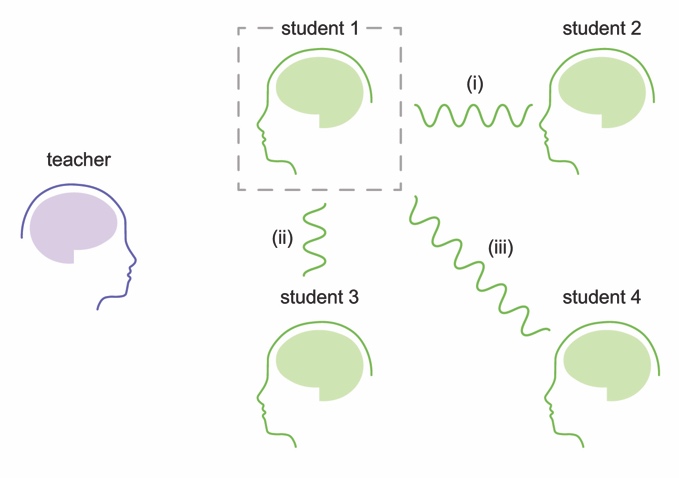

**Figure S1**: Calculation of brain-to-brain synchrony

Brain-to-brain synchrony (CCorr) values were computed between each student and all other students in the group as well as between each student and the teacher.

**Figure S2**: The relationship between brain-to-brain synchrony and delayed retention across three different frequency bands: Theta (4-7Hz; A), Alpha (8-12Hz; B), and Beta (13-20Hz; C). Only alpha-band brain-to-brain synchrony significantly predicted delayed retention (p<0.05).

**Figure S3**: The relationship between brain-to-brain synchrony and delayed retention across three different regions of interest: Posterior (A), Central (B), and Frontal (C).

**Table S1.** Subset of questions presented on pre-, post-, and delayed post assessments.

| Lecture | Example question | Example answers |
| --- | --- | --- |
| Bipedalism | Which of the following is a negative consequence of using bipedal locomotion? | A) Tarsal tunnel syndrome  B) Slipped discs  C) More frequent breaks in the femur  D) Constipation |
| Insulin | Insulin is produced by what type of cells? | A) Beta cells  B) Alpha cells  C) T cells  D) Red blood cells |
| Niches and Habitats | Each species occupies a unique niche because…. | A) Species are so varied  B) Resources are limited in a habitat  C) There are a limited number of niches  D) There is a limit to the size of a niche |
| Lipids | Which of the following is not a solid at room temperature: | A) Unsaturated fats  B) Saturated fats  C) Trans-fats  D) Phospholipids |

**Table S2.** Summary of EEG channels, ICA components and epochs that were rejected during preprocessing. Numbers reflect mean ± s.d.

|  | Student Datasets  (N=123) | Teacher Datasets  (N=34) |
| --- | --- | --- |
| Number of rejected channels | 2.05 ± 2.25 | 4.00 ± 2.10 |
| Number of rejected ICA components | 1.65 ± 0.50 | 1.91 ± 0.38 |
| % epoch rejected | 11.34 ± 6.64 | 27.41 ± 12.47 |
